# A synthesis of the evidence for the acoustic space partitioning hypothesis

**DOI:** 10.64898/2026.04.23.718589

**Authors:** Katherine González, Brian C. Leavell, Jeffrey R. Lucas, Ximena E. Bernal

## Abstract

Acoustic interference is a critical factor driving the evolution of communication systems. In mixed-species choruses, competition for acoustic space is expected to drive signal differentiation among heterospecific signals. The acoustic space partitioning hypothesis proposes that species differentiate their signals to reduce overlap and thereby reduce acoustic interference. Despite ongoing debates in niche theory, studies in animal communication have remained disconnected from these conversations, and no critical evaluation of this hypothesis has been conducted. We performed a systematic review to assess empirical support for acoustic space partitioning and evaluate the conceptual and methodological approaches used to test it. We found that two-thirds of studies conclude that the acoustic space is partitioned, albeit with a strong taxonomic bias toward anurans. However, studies rarely account for key assumptions of the hypothesis, including co-signaling, limited acoustic space, and signal masking at the receiver. Without explicit evidence of conditions for acoustic interference, signal differentiation alone is insufficient to infer that competition is driving partitioning, since this outcome may also arise from alternative processes. By integrating sensory ecology, we provide a framework to reconcile signal-structure differentiation with receiver perception, thereby improving our understanding of how communication systems evolve in mixed-species choruses.

## I. INTRODUCTION

Acoustic interference is a common challenge for signalers in mixed-species choruses [1,2]. These signaling environments erode relevant properties of communicative signals and, in the case of mating signals, lead to loss of mating opportunities, pairing with unpreferred conspecifics, or, worse, heterospecifics [3,4]. Acoustic interference arises when signals overlap in space and time, affecting signal detection or discrimination. This process reduces the distance at which signal amplitude remains above the receiver’s detection threshold (hereafter, the active space [5]). Given the high costs of missed mating opportunities and lower predator detection associated with acoustic interference [4,6,7], behavioral and auditory strategies have evolved to overcome this communication challenge. These strategies have attracted attention across taxonomic groups [8], as they provide a foundation for understanding the mechanisms underlying signal evolution.

One mechanism commonly invoked to explain heterospecific signaling in choruses (hereafter, co-signalers [9]) is signal differentiation to avoid acoustic interference. Differentiation can occur along three dimensions of the acoustic niche: (*i*) acoustic space: acoustic features of the signals of species in the aggregation [10–12], (*ii*) spatial dimension: signaling from different habitats to segregate across physical space [13,14], or (*iii*) temporal dimension: signaling at different times of the day [15] or year [16] (figure 1A). However, in mixed-species choruses, only the acoustic space can be partitioned, as signalers overlap in time and space (figure 1B1, B2). The acoustic space partitioning hypothesis thus proposes that, in mixed-species choruses, acoustic interference is reduced through signal differentiation, thereby decreasing detection and discrimination errors by receivers. A prediction extending from this hypothesis is that species use unique communication channels, similar to how radio stations broadcast messages at different radio frequencies [17,18]. Although this hypothesis is commonly invoked to explain signal differences among co-signalers, its explanatory power remains untested despite concerns about current evidence [19–21].

**Figure 1.**
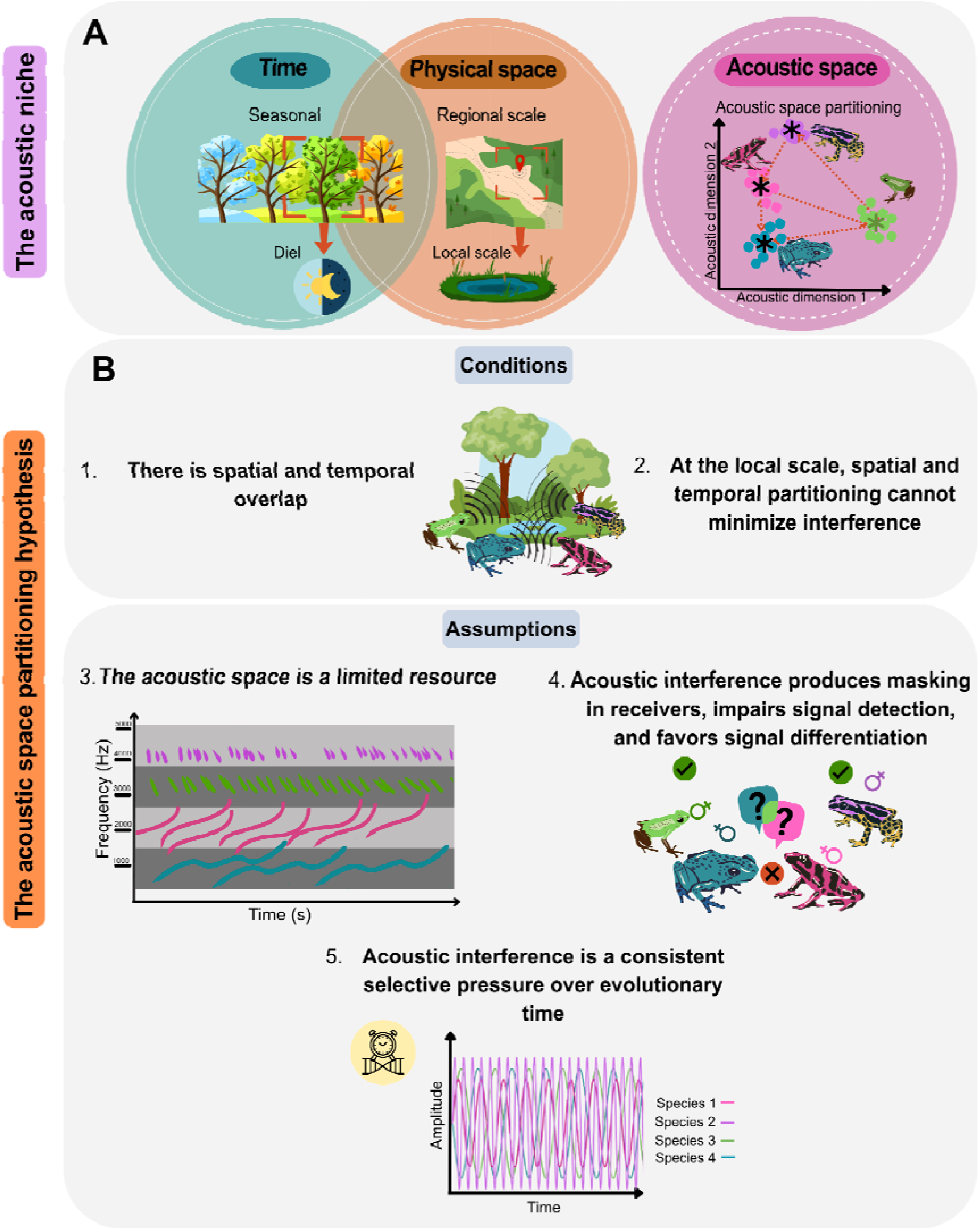
Components of the acoustic *niche* and acoustic *space* that lead to partitioning. A. The acoustic *niche* comprises three dimensions along which signals can differentiate between species: time (when to signal), physical space (where to signal), and signal properties (*acoustic space*). Definitions of key conceptual terms can be found in the glossary (Sup. Mat). B. Visual representation of the conditions (1-2), and assumptions (3-5) of the acoustic space partitioning hypothesis.

Unlike frameworks emphasizing environmental conditions, sexual selection, or evolutionary drift as drivers of signal divergence and speciation [22], bioacoustics has framed partitioning of the acoustic space through niche theory around reducing acoustic interference. In species-rich aggregations, the acoustic space is expected to be saturated, thereby limiting the coexistence of species sharing similar signal traits [10,18] (figure 1B3). From an evolutionary perspective, receivers can incur errors imposed by difficulty in detecting or decoding mating signals in the presence of heterospecific signals. These mating errors are expected when overlapping signals cause acoustic interference that cannot be alleviated by temporal or spatial segregation (figure 1B4). Furthermore, selective pressure imposed by acoustic interference must be experienced consistently and predictably over evolutionary time to shape both signal differentiation and the receiver’s ability to perceptually segregate heterospecific signals from conspecifics (figure 1B5) [4]. Thus, signal partitioning is likely to occur in systems with persistent temporal and spatial overlap among species, such as recurring breeding choruses. Such partitioning arises from differences in signal design due to evolutionary divergence, either through changes in fixed traits or evolved plasticity [23]. Together, these processes delimit acoustic space partitioning as a conditional outcome of competition for acoustic space, yet the extent to which this hypothesis is supported remains unknown.

In this review, we synthesize studies examining the acoustic space partitioning hypothesis to evaluate evidence for signal differentiation as the outcome of acoustic interference in mixed-species choruses. To evaluate the current state of this hypothesis, we systematically reviewed the literature across taxonomic groups, characterizing patterns in the taxa studied, the acoustic features evaluated, and the methodological approaches used, thereby revealing general trends and biases. We synthesize the main findings regarding the conditions under which acoustic space partitioning is evaluated and discuss conceptual and methodological challenges obscuring the role of acoustic interference in signal evolution.

We then focus our discussion on acoustic space partitioning in anurans, as these organisms provide an ideal opportunity to examine this hypothesis, given that their signaling ecology closely aligns with the conditions required for partitioning [19]. Frogs commonly breed in dense choruses within restricted areas and during a limited time, restricting spatial or temporal partitioning. Additionally, most frog acoustic signals are stereotypic [24,25] and acoustic interference exerts strong selective pressure on their communication system because receiver errors caused by masking interference incur substantial fitness costs [26]. Given the critical role that sensory processing plays in mediating signal detection and minimizing acoustic interference, we integrate studies of anurans’ auditory systems into our discussion.

## II. METHODS

We systematically reviewed studies investigating the acoustic space partitioning hypothesis of mating signals in mixed-species choruses. Literature searches were performed in Scopus and Google Scholar using predefined search strings (Supplementary material, figure S1). Searches included all document types, all years, and all languages to minimize language-related bias [29]. The initial search, conducted between May 2023 and January 2024, identified 31 studies. We then expanded the search to include 64 Scopus and 136 Google Scholar studies, of which only 35 (15%) matched the goals for this review. Using this initial set of studies, we conducted backward and forward reference searches using Citationchaser [30], which identified an additional 4,713 records. After removing duplicates and screening abstracts, 262 papers partially met our criteria. Following a full-test assessment, 78 studies were retained for synthesis (Supplementary material, figure S1).

We excluded non-peer-reviewed records, including books, theses, conference abstracts, and inaccessible articles. Given our interest in understanding partitioning in mating contexts, we excluded two studies evaluating non-mating signals. We included studies assessing spatial or temporal (diel or seasonal) partitioning only when acoustic features were also evaluated. We further restricted inclusion to studies examining airborne signals as the primary sensory modality, excluding chemical, vibrational, and waterborne communication. To examine the evidence of partitioning in mixed-species choruses, only studies including two or more species were retained.

From each study, we retrieved conceptual and methodological information describing how acoustic space partitioning was evaluated. Specifically, we recorded the taxonomic group studied, number of species included, acoustic features measured, and whether the acoustic space was explicitly defined. We assessed reported support for acoustic space partitioning based on the authors stated conclusions. We also recorded whether studies explicitly assessed the role of plasticity in acoustic space partitioning.

We further synthesize the methodological approaches used to quantify partitioning, including whether signal distributions in acoustic space were compared against null models, how acoustic features were quantified, and whether multicollinearity among parameters was evaluated. When available, we also extracted information on assemblage characteristics, including the number of species per aggregation and the spatial scale of the study. Due to inconsistencies in how spatial scale and assemblage composition were reported across studies, these data were summarized descriptively (Supplementary material, figure S1).

To assess the association between study factors and the outcome of support, we conducted various analyses with the outcome as the response variable. We performed our analyses in R Core Team (2022) using RStudio Team (2021) [31]. We used chi-square tests to examine relationships between categorical variables, including taxonomic group, use of null models, and reported support for partitioning. We fit logistic regression models using the logit function to assess the effects of continuous variables, such as the number of features and species, on the study outcome. Model fit was evaluated using Nagelkerke’s R^2^ values [32]. All studies included in the review and data retrieved are provided in the supplementary material.

## III. RESULTS

### III. A. General trends in the study of acoustic space partitioning across taxa

#### Chronological and taxonomic patterns in studies

The first study investigating acoustic space partitioning was published in the late 1950’s [10] and was followed by a few studies examining this phenomenon in subsequent decades [1,33–35]. The number of studies increased dramatically between 2000 and 2023, with 81% of studies published during that period (figure 2). Despite the increasing number of studies, however, research efforts across taxa remain relatively uneven. Anurans are the most studied taxonomic group, comprising more than half of the literature (N = 42 studies), followed by birds (N = 21 studies) and insects (N = 14 studies) (figure 2). Studies on marine mammals are rare, with only three studies addressing partitioning of mating signals. Across studies, there is no association between the taxonomic group investigated and the reported support for the hypothesis (χ² = 1.72, p-value > 0.05). Lastly, only three studies included multiple taxa, such as insects, birds, and anurans [36] or insects and birds [37,38].

**Figure 2.**
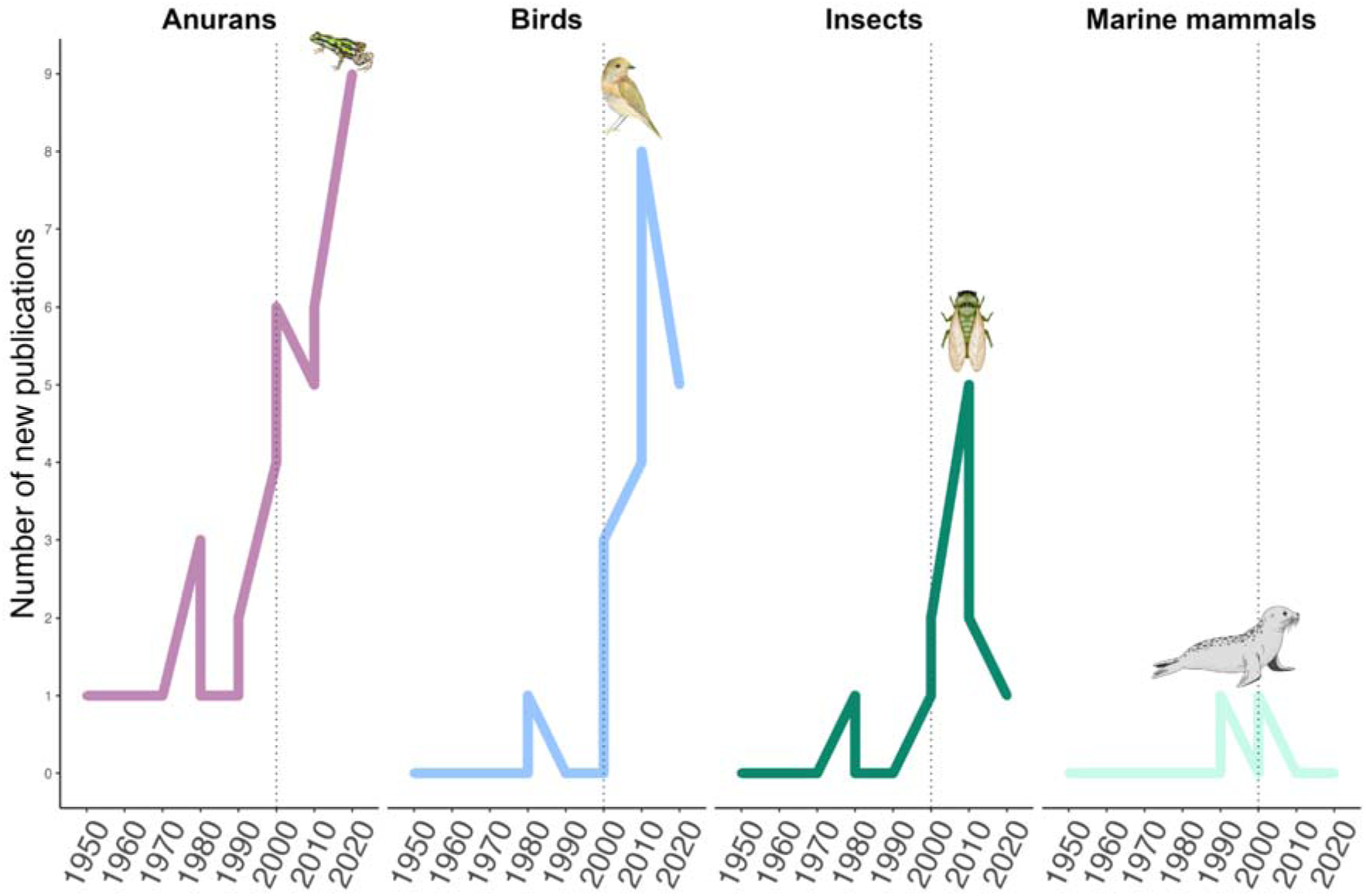
Chronological and taxonomic characterization of studies investigating the acoustic space partitioning hypothesis. Number of studies published per decade since the first publication in 1958 through 2023. The dotted vertical line marks a surge in studies at the turn of the century.

#### Niche dimensions used to assess partitioning of acoustic space

Of the 78 studies meeting the inclusion criteria, 25% examined the acoustic space exclusively whereas 74% assessed differentiation in at least one additional dimension of the acoustic niche, either temporal [39–41], spatial [13,42,43], or a combination of those and signal features [1,44,45]. One study did not clearly report an outcome for signal differentiation because assessing this hypothesis was not aligned with their goals [46].

#### Definition of the acoustic space

Only 12% of the studies explicitly defined what the authors meant by “acoustic space”, but in all cases, studies referred to the signal structure of the aggregation (Supplementary material, table S1). When defining the acoustic space, most studies (8 out of 10) included terms such as “limited” or “resource,” aligning with early ideas framing the acoustic space as an ecological resource susceptible to competition [18]. While some recent studies continue to adopt this resource-based definition [47,48], others have decoupled the acoustic space from being an ecological resource per se by not framing it as a product of competition [21,49]. Instead, some studies describe the acoustic space as a physical continuum of an infinite number of sound properties, without explicitly referencing the potential for resource limitation (e.g. [21,50]). Nonetheless, as we continue assessing the evidence for this hypothesis, and to avoid presupposing any underlying mechanisms, we followed an approximation to the definition proposed by Chhaya et al. (2021) (Glossary, Supplementary materials).

#### Acoustic features used to characterize the acoustic space

Consistent with variation in the concept of acoustic space, studies also differed in the features used to characterize signal structure. Across all studies included here, 100 distinct features were used to characterize signals, with a median of 5 per study. Studies of anuran calls used the highest feature diversity (62 features in total; mean ± SD = 5.85 ± 3.36 features per study), followed by bird studies (46 features; mean ± SD = 5.57 ± 4.22 features per study). In insects and marine mammals, studies used fewer features (insects: 35 features, mean ± SD = 4.6 ± 3.6; marine mammals: 7 features, mean ± SD = 3 ± 1.41). The use of features to describe the acoustic space was inconsistent across studies: 68% of all features appeared only once or twice, and among these, 60% were not explicitly defined by researchers.

Dominant frequency was the most commonly used acoustic feature, reported in 71% of studies, followed by frequency bandwidth (35%). This pattern was consistent across taxonomic groups (figure S2). For example, dominant frequency was quantified in 71% (30 out of 42) of anuran studies, whereas frequency bandwidth was evaluated in 38% of the cases. Among temporal properties, call duration was included in half of the studies, while other features such as the number of notes, note duration, and pulse rate were measured less frequently (∼20% of studies; figure S2). Although correlations among features are common, only 20% of studies (17 of 85) explicitly evaluated collinearity among traits.

### III. B. Reported support for acoustic space partitioning

After examining the reported outcomes, we found that 48 out of 78 studies (62%) claimed to have found evidence of support for the hypothesis, whereas 26 (33%) reported no support. Reported outcomes were not associated with publication year (χ^2^ = 5.98, p > 0.05), taxonomic group (figure 3; χ^2^ = 1.72, p > 0.05), or number of features used (z = −0.109, s.e. = 0.067, p-value > 0.05). However, reported outcomes varied according to the niche dimensions examined. In studies focusing on mixed-species choruses, and thus assessed acoustic space exclusively, 90% (18 out of 20) claimed consistent supporting evidence for acoustic space partitioning. In contrast, when studies evaluated acoustic space alongside temporal and/or spatial dimensions, the reported evidence supporting partitioning was lower (64% of cases). One study reported mixed support, finding partitioning in 4 of the 11 assemblages examined [19].

**Figure 3.**
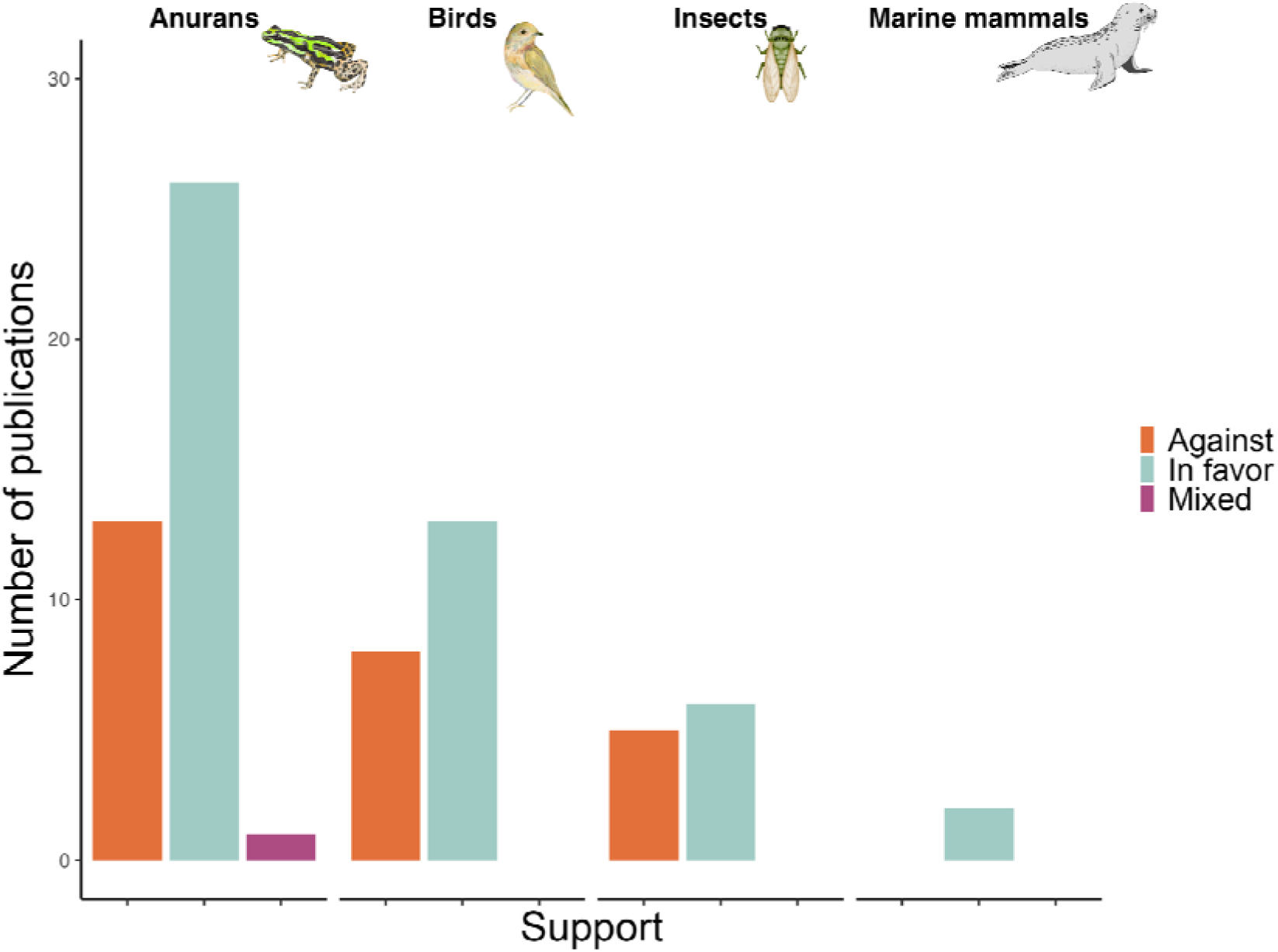
Purported evidence for the acoustic space partition hypothesis across taxonomic groups. Anura is the most studied group, with 2 out of 3 studies supporting acoustic space partitioning (in favor = 26, against = 13, mixed = 1).

### III. C. Assessment of the evidence on acoustic space partitioning

#### Evidence for co-signaling in mixed-species choruses

A central assumption for acoustic space partitioning is that heterospecific signals result in acoustic interference within mixed-species choruses (figure 1). Thus, studies are expected to provide information on the spatial and temporal conditions of the aggregation. For instance, studies could explicitly describe whether signals were recorded simultaneously, whether individuals signaled in close proximity to one another, or whether species consistently overlapped during breeding events. We found, however, that studies vary widely in how they define and characterize focal mixed-species choruses.

Most studies (65%) reported focusing on a single mixed-species chorus, whereas others assessed multiple aggregations across broader spatial scales (Supplementary material, table S2). Terminology used to define aggregations was inconsistent, with studies referring variously to sites, locations, habitats (e.g. water bodies or forests), regions, or geographic areas, often without specifying whether species were syntopic or actively co-signaling. It is thus unclear whether species included in the same assemblage overlapped at the spatial scale required for acoustic interference. Indeed, direct evidence of heterospecific co-signaling was not clearly reported. Therefore, the spatial and temporal contexts of signal production by species included in a given study may not overlap vocally.

Given that the number of species within an aggregation is used as a proxy for how packed the acoustic space is [19], we examined the role of aggregation size in signal partitioning. We found great variation in the number of species per focal aggregation, ranging from two to several dozen (e.g. 85 species in a bird assemblage [51]), with a median of 12 species per aggregation. Although studies on birds and insects typically included more species per aggregation (median = 17 and 12, respectively) than in anurans or marine mammals (median = 10 and 3, respectively), the number of species included was not associated with reported support for the hypothesis (Z = −0.164, p-value > 0.05). Reporting of species composition was also inconsistent, with 31% of studies pooling signals from species across multiple assemblages without specifying species composition at the level of individual aggregation (e.g. 300 species distributed across 90 assemblages [20]).

Lastly, we assessed whether studies evaluated temporal overlap among signalers at scales relevant to acoustic interference. Although signal differentiation at fine temporal scales (e.g., within seconds/minutes) is expected to be more relevant for masking release, studies reported coarse temporal overlap (season/day). Studies typically described when sampling occurred (e.g. seasons, diel periods, months), without addressing how consistently co-signalers experience acoustic interference. Because fewer than a third of studies directly assessed both spatial and temporal dimensions in addition to acoustic space (24 out of 78), it is unclear whether segregation across space or time reduced acoustic interference.

#### Quantification of the acoustic space: implications for competition

A third of the studies investigating acoustic space partitioning relied on descriptive approaches, characterizing signal spectral and temporal features through visual inspection of spectrograms and basic statistics (Supplementary material, table S3a). In these studies, authors examined whether signals differ among species, an outcome often interpreted as evidence of past competition. Because such assessments are typically based on summary statistics, they provide detailed descriptions of signal spectral and temporal features. Although these analyses are ideal for data exploration, they are limited when evaluating how signal overlap may result in acoustic interference. In some cases, descriptive approaches were complemented by additional analyses (13 studies), such as dimensionality reduction methods (e.g., principal component analysis and discriminant function analysis), to identify features contributing to signal differences [52].

A subset of studies evaluated acoustic space partitioning by measuring the degree of dispersion among signals of different species within a multidimensional acoustic space. These approaches, derived from community ecology, interpret trait overdispersion as evidence of competition for a limited resource [53]. Within this framework, the implicit rationale is that if acoustic space is structured by competition, species’ signals should be more widely dispersed than expected by chance. Signal overdispersion is usually calculated using the distances between species’ signals in acoustic space throughout Cartesian coordinates (e.g. Euclidean distances), or distance-based clusters (e.g., nearest-neighbor analysis) (Supplementary material, figure S1, table S3). Acoustic features are combined so that increases in signal differences result in proportional increases in calculated distances [54]. Although comparisons with null expectations for random distributions are central to distinguishing patterns shaped by competition from chance [19,55], this approach remains rare in studies of acoustic space partitioning (2 of 15 studies).

Rather than testing overdispersion, the most common approach used quantifies overlap among species’ signals in acoustic space. Approximately 44% of studies compared distributions of signal traits across species, testing whether they derive from similar distributions or exhibit minimal overlap (e.g. Kolmogorov-Smirnov test; Supplementary material, table S3). A smaller proportion of studies (12%) discretized the acoustic space into categories defined by features (e.g., frequency bands or pulse-rate intervals) and quantified overlap by counting the number of categories shared between species pairs. For example, one anuran study divided the total call frequency range into 200Hz bands, estimating overlap as the number of bands shared between species pairs [56]. Similarly, category-based approaches use diversity indices (e.g., inverse Simpson or Czekanowski indices) to compare the distribution of species signals across feature categories, whereas other studies used predictive statistical models (e.g., Pearson’s correlation, LMM, GLMM; Supplementary material, table S3) to test whether species identity explains signal structure. Lastly, a small number of studies (20%) combined multiple analytical approaches, typically using exploratory methods to reduce the dimensionality of signal structure before evaluating signal differentiation.

#### The role of plasticity in partitioning of the acoustic space

Signal differences among species may reflect fixed or short-term plastic responses. Plasticity can arise as an adaptive response under variable co-signaling conditions, such as fluctuating signaler abundance, favoring plastic signal adjustments to reduce acoustic interference. We found, however, that most assessments of acoustic space partitioning relied on signal measurements pooled from a single recording context. Only two studies tested whether features were context-dependent by experimentally manipulating the acoustic environment [57,58]. In both cases, researchers presented frogs with heterospecific signals (invasive species calls) and quantified their vocal responses to assess differences in signal traits. Although changes in signal spectral and temporal traits are species-specific, in some cases, frogs increased the spectral overlap between their calls and those of the invasive species [58]. While lack of a shared evolutionary history between these species confounds the interpretation of such findings in the context of signal partitioning, they highlight the relevance of considering plastic, heterospecific vocal interactions, which may modulate acoustic interference. Incorporating context-dependent approaches provides a promising path for linking acoustic interference to short-term responses and long-term signal differentiation.

#### Receiver perception and acoustic interference

Nearly all studies inferred acoustic interference from patterns of signal differentiation or overlap in acoustic space from the signaler’s perspective, with two studies explicitly incorporating receiver responses using behavioral and neurophysiological approaches [11,12]. One of these studies used playback of calls of sympatric populations of *Hyla cinerea* and *H. andersonii* to examine female *H. andersonii* responses to conspecific and heterospecific calls [11]. While this study met the inclusion criteria for this review, it does not directly examine acoustic interference on female responses, as these species reduce opportunities for acoustic interference by calling from different microhabitats. Such spatial partitioning has likely reduced selection for signal differentiation. Ultimately, this study provides limited insight into the role of receiver responses in mediating the effects of acoustic interference on signal differentiation.

The second study directly examined perceptual segregation of signals in three co-signaling cicada species (*Cicada orni*, *Cicadatra atra*, and *Lyristes plebejus*). These species produce signals with spectral differences, and the ears of males are tuned to the frequencies of their conspecific song. The tympanal membranes of females, however, are broadly tuned to male signals. Although the cicadas’ signals matched the prediction of signal differentiation in acoustic space, those differences do not translate to apparent perceptual separation for one target receiver. Instead, females cope with acoustic interference through mechanical and neuronal processing beyond the peripheral auditory system [12], showing that perceptual discrimination can emerge through mechanisms not evident from peripheral tuning alone. Thus, signal differentiation may not necessarily correspond to reduced perceptual overlap for receivers. While signal differences are commonly interpreted as evidence of reduced acoustic interference, in the absence of receiver-based evaluation, such conjecture may be unsupported.

**Box 1.**
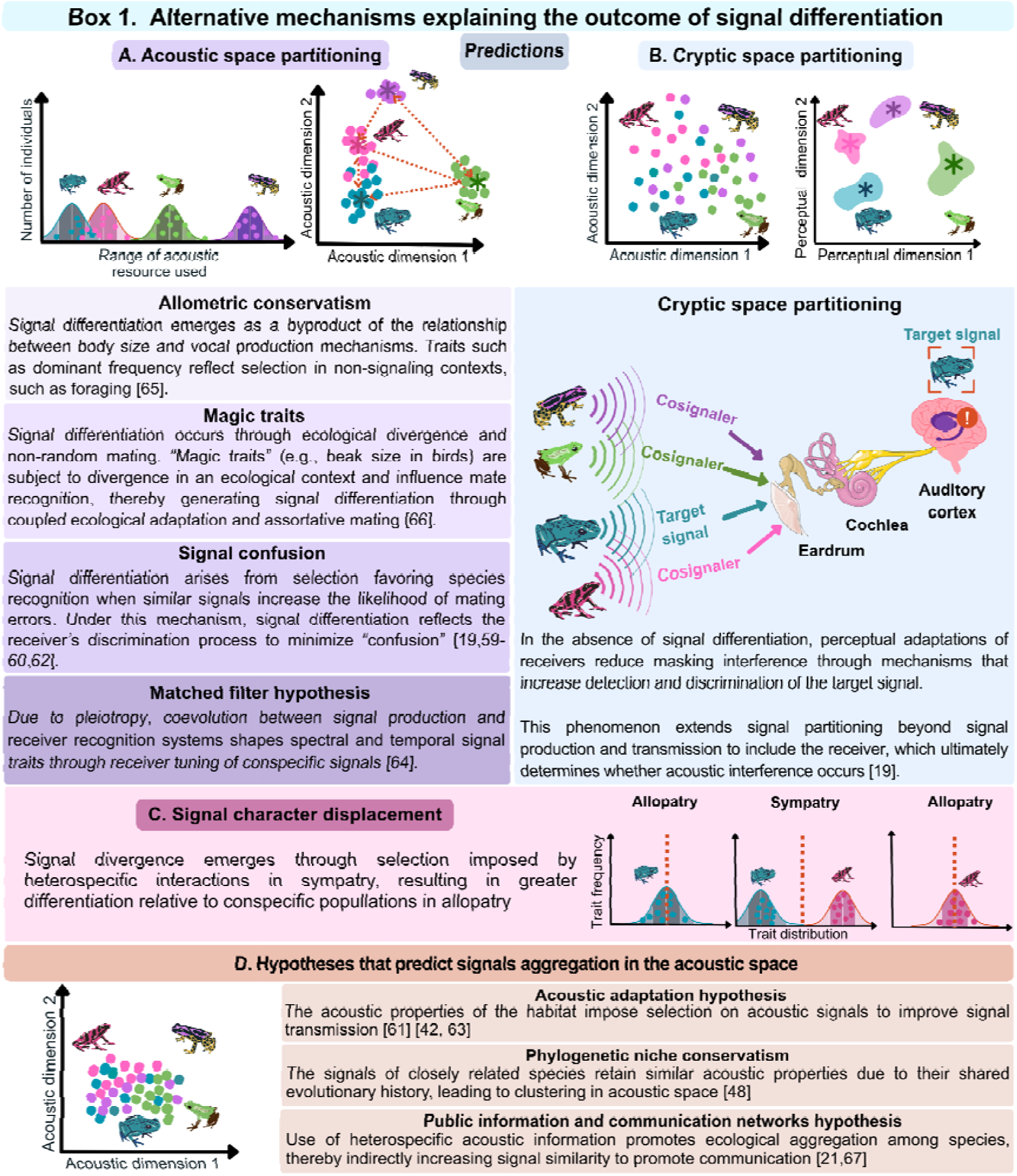
Alternative processes, additional to acoustic interference, can lead to signal differentiation. Considering the signaler, at least four other mechanisms explain segregation of the acoustic space (A). Taking the receiver into account (B), cryptic sensory processing can decouple signal interference. Signal character displacement can provide further evidence of the role of acoustic interference (C). Other hypotheses predict convergence of signals in acoustic space, resulting in aggregated rather than segregated acoustic space (D).

## IV. DISCUSSION

The acoustic space partitioning hypothesis has long been invoked to explain signal differentiation in mixed-species choruses. We show that empirical support is generally consistent, with approximately two out of three studies reporting evidence of acoustic space partitioning. Despite conceptual and methodological variation across studies, the prevalence of support across taxonomic groups suggests signal differentiation is widespread. However, interpreting such a pattern as evidence of competition driven by acoustic interference remains difficult. In many cases, studies infer signal partitioning without evaluating the extent to which acoustic space is limited, or whether observed differences reduce masking by the receiver.

### IV. A. Evidence for the acoustic space partitioning hypothesis: General lessons

Acoustic interference, the selective force invoked by the acoustic space hypothesis, requires sustained temporal and spatial overlap among heterospecific signalers affecting signal transmission and reception. However, many studies infer potential for acoustic interference from broad-scale sympatry or regional co-occurrence, despite evidence that co-occurrence alone does not necessarily result in ecological interactions or co-signaling [68]. Species may co-occur at a given habitat or region, yet segregate their signaling activity across time or space, reducing opportunities for acoustic interference [39,41,69]. When overlap along these axes is not evaluated, there may be a mismatch between the scale at which signaling occurs and the scale at which acoustic space partitioning is inferred. As a result, signal differentiation may be attributed to acoustic partitioning even when competition for acoustic space is unlikely.

Even when the ecological conditions for acoustic interference are met, studies differ substantially in how they quantify and interpret acoustic space partitioning, thereby contributing to variation in reported support. Early studies relied largely on descriptive statistics and visual inspection of spectrograms, with assessments of signal differentiation relying heavily on researcher judgment. Such analyses can thus lead to biased conclusions about whether signals from different species are sufficiently distinct to avoid acoustic interference. More recently, studies have used several quantitative approaches, including overlap metrics, distance-based measures, and multivariate analyses. These approaches differ fundamentally in their biological assumptions and the aspects of signal variation they emphasize. Distance-based methods, for example, typically rely on the central tendency of a species’ acoustic feature (e.g., the centroid) and further reduce signal dimensionality using techniques such as principal component analysis. While useful for summarizing complex, multivariate acoustic data, these approaches can obscure intraspecific signal variation and may be sensitive to limited or uneven sampling, potentially inflating interspecific distances [70,71].

Although distance-based approaches are used for their ability to summarize complex data, robust inference of signal segregation requires evaluating observed patterns against null expectations. Chek et al. (2003) examined the consequences of omitting such comparisons by revisiting studies reporting acoustic space partitioning in anuran assemblages. Their findings show that when signal distributions are compared with random expectations, few assemblages (4 of 11) exhibit statistically significant overdispersion [19]. Although null models are central to distinguishing competition-driven patterns such as overdispersion, we found they are rarely implemented, increasing the likelihood of false positives. Thus, the relatively high proportion of studies we found reporting support for acoustic space partitioning may be inflated, as researchers infrequently excluded whether such patterns arose by chance.

Regardless of whether overdispersion is detected, it remains unclear if species are competing for a limited acoustic resource. Although acoustic space partitioning is based on niche competition theory [18], few studies explicitly define acoustic space, and when they do, resource availability is assumed rather than tested. Because the acoustic space is inherently difficult to quantify, studies have estimated resource availability indirectly by assessing acoustic saturation. In general, mixed-species choruses with wider niches, high species richness, and limited acoustic space are expected to yield saturation patterns promoting acoustic partitioning [19]. For example, insect richness can drive birds’ and frogs’ use of acoustic space in tropical soundscapes [36]. However, saturation is also likely associated with environmental conditions modifying the acoustic space available. Natural noise, such as running water or rainfall, may constrain acoustic space availability, limiting opportunities for signal differentiation [72]. Overall, the challenge of delimiting the acoustic space as a resource highlights important differences between how competition is evaluated in ecology and how it is inferred in animal communication studies.

An additional consideration, beyond the methodological approaches, is that several alternative mechanisms can generate signal differentiation, making this pattern a non-exclusive prediction of the acoustic space partitioning hypothesis (Box 1A). In frogs, for example, dominant frequency is correlated with vocal fold length, a trait predicted by body size [73]. Because frogs are gape-limited predators, ecological factors, such as diet, may indirectly shape call frequency [74] (but see [75,76]). Since similar relationships between body size and dominant frequency occur in other groups (birds: [77], insects: [78,79]), divergence in dominant frequency resulting from selection imposed in non-signaling contexts likely extends beyond anurans (Box 1A). One study in our review, however, examined acoustic space partitioning alongside allometric conservatism. This study investigated two sympatric frog species with similar size and body proportions, and, despite similar diets, their signals differed in dominant frequency and call duration [80], showing that signal divergence can emerge independently of ecological divergence. Further work is needed to better understand how selection imposed on body size in non-signaling contexts modulates acoustic interference. Habitat characteristics, for instance, can indirectly influence signal properties through putative “magic traits” (Box 1A, [81]), offering an alternative explanation for signal divergence patterns (e.g. [66]).

Similarly, other processes involving receiver-mediated mechanisms, including signal confusion and matched-filter mechanisms (Box 1A), can also result in patterns of signal differentiation. While signal divergences can result from diverse processes, failure to detect this pattern does not necessarily reject the idea that acoustic interference has imposed selection shaping species-specific communication channels. Perceptual adaptations in receivers may facilitate signal detection and discrimination despite substantial overlap in acoustic space (cryptic space partitioning: Box 1B).

As is well recognized, detecting the signature of natural selection from past interspecific competition in shaping the niches of existing species when competition is not currently occurring (ghost of competition past; see [82]) can be difficult. Character displacement has been the primary approach for examining historical competition in ecology [83], and analogous approaches targeting vocalizations can be used (signal character displacement, Box 1C). When considering mating signals, however, selection to increase reproductive isolation (i.e. reinforcement, [84]) may also shape signal divergence. To further complicate the challenge, the multiple hypotheses predicting signal differentiation are not mutually exclusive, and different selective pressures could act in conjunction with acoustic interference to drive signal divergence. Yet there are competing hypotheses, such as the acoustic adaptation hypothesis and phylogenetic conservatism, that predict distinct patterns of signal clustering in acoustic space (Box 1D).

### IV. B. Anuran choruses: a case study examining the role of acoustic interference

Signal differentiation among co-occurring species is frequently interpreted as evidence that acoustic interference has diminished through acoustic space partitioning. However, as outlined above, differentiation in signal structure alone is insufficient to infer reduction in interference. In frog choruses, numerous studies document signal differentiation in one or more signal traits among co-signaling species, most commonly in dominant frequency or call duration (e.g. [10,11,33]). Given that acoustic interference occurs when overlapping signals compete for neural representation and perceptual salience [85,86], signals that appear well separated in acoustic space may still result in acoustic interference by the receiver. Similarly, signals with overlapping spectral features may remain discriminable due to narrow filtering or temporal coding mechanisms.

Experimental work in anurans illustrates the potential mismatch between signal differentiation and functional interference by the receiver. Considering auditory perception, it is well established that close match between signal frequency components and auditory processing mechanisms, such as peripheral auditory filters, can minimize masking interference [85,87–90]. Nevertheless, frog receivers often discriminate conspecific calls in dense choruses despite substantial spectral overlap with heterospecific signals. Such discrimination is mediated by species-specific auditory filters and sensitivity to fine-scale temporal features enhancing masking release [85,88–90]. Beyond peripheral tuning, anuran receivers can also employ additional adaptations that reduce the impact of background noise. For example, resonance properties of the lungs in American green tree frogs (*Dryophytes cinerea*) increase spectral contrast with heterospecific signals by attenuating signal energy outside of frequency peaks of the conspecific call [91].

Although studies of acoustic space partitioning have focused on spectral differentiation, temporal acoustic components represent an underappreciated yet critical axis of communication. Species identity can be encoded through fine spectro-temporal structure and signal envelope features, such as rapid frequency or amplitude modulation [92,93]. In frogs, even when signals overlap spectrally, hearing adaptations as early as in the periphery extract species-specific temporal features, facilitating the processing of temporal signals [94]. For example, females of Cope’s tree frogs (*Dryophytes chrysoscelis*), rely primarily on pulse rate for species recognition, whereas gray tree frog (*Dryophytes versicolor*) females integrate inter-pulse interval, absolute pulse rate, and pulse rise time [95,96]. Similar temporal discrimination mechanisms occur across taxonomic groups, including insects [97], frogs [98,99], and birds [100,101]. As a result, cryptic acoustic partitioning may arise at the receiver even when signal overlap may exist in the acoustic space (Box 1B).

Together, these lines of evidence underscore that reductions in acoustic interference arise not solely from signal differentiation but also from receiver signal processing. Signal differentiation alone provides, at best, indirect support for the hypothesis unless it is explicitly linked to reduced auditory masking or improved discrimination ability by receivers. Receiver perceptual space is arguably a neglected, but critical, fourth dimension of the acoustic niche.

### IV. C. Ecological conditions for acoustic space partitioning

The acoustic space partitioning hypothesis assumes that acoustic interference imposed sustained selection on signal structure over evolutionary time. However, even when heterospecific signalers overlap in space and time, selection arising from acoustic interference may be transient or weak, favoring short-term plastic responses rather than long-term divergence. Under these conditions, the extent to which acoustic interference contributes to signal differentiation may depend on whether signal traits are evolutionarily constrained or sufficiently plastic to respond to changing acoustic environments. Studies of mixed-species aggregations are beginning to clarify the ecological conditions under which acoustic interference generates patterns of signal differentiation regardless of competition.

Recent work is increasingly examining signal variation across ecological contexts and the role of plastic responses in acoustic space partitioning. Short-term shifts in dominant frequency can occur in response to natural (e.g., streams, [102]) and anthropogenic noise [103]. Similar plastic responses may also arise under novel competitive conditions created by invasive species, which can alter the acoustic environment (reviewed in [104]). For example, native frogs can deploy plastic shifts in call frequency in response to calls of invasive species [57,58,105]. Nonetheless, it is unclear whether such plasticity reduces masking [104].

The temporal components of anuran calls are particularly plastic in response to social and acoustic contexts [3,106]. Frogs can exploit silent gaps to reduce overlap with other signalers [107], effectively partitioning the acoustic niche along a micro-temporal axis (e.g. seconds or minutes). Such fine temporal adjustments can occur in response to artificial noise, with Neotropical treefrogs (*Boana bischoffi* and *B. leptolineata*) exposed to traffic noise altering call rate or call duration depending on noise intensity and species identity [108]. Similar adjustments take place in response to novel signalers, such as invasive species [57,109]. Plasticity in temporal traits may first emerge in response to acoustic interference as a precursor of long-term signal divergence.

Because the intensity and predictability of acoustic interference vary across ecological contexts, plastic signal adjustments may be particularly advantageous under dynamic signaling conditions. Spatiotemporal variability in ambient noise alters the intensity and predictability of acoustic interference, creating a dynamic selective landscape for co-signaling species. From a coexistence perspective, the probability of masking should depend not only on signal overlap but also on the relative abundance of co-signaling species. In simulated insect choruses, effective acoustic overlap (representing the highest level of masking likely experienced by receivers) was greater in choruses with high species evenness relative to those with a few dominant species [110]. Similarly, in frogs, shifts in call frequency by male *Eleutherodactylus coqui* and *E. portoricensis* occur only when both species are present at similar densities, suggesting that acoustic interference depends on relative signaler density [111]. Although signal structure remains an important factor in acoustic interference, variation in signaler abundance introduces a density-dependent component that can modulate both the strength and symmetry of masking in mixed-species choruses.

## V. CONCLUSIONS AND FUTURE RECOMMENDATIONS

The acoustic space partitioning hypothesis has been widely invoked to explain signal differentiation in mixed-species choruses. Although about two-thirds of the studies reviewed reported evidence consistent with this hypothesis, this result should be interpreted with caution, as the main prediction tested is signal differentiation, and this outcome can arise through mechanisms other than acoustic interference. Moreover, signal differentiation is often interpreted as resulting from past competition for acoustic space, without evaluating interspecific competition imposed by acoustic interference. Acoustic interference may instead be solved through receiver adaptations, signal plasticity, and context-dependent signaling strategies. To improve inference and facilitate direct assessment of the hypothesis, future work would benefit from integrating ecological, perceptual, and evolutionary perspectives on acoustic interference. In particular, future studies should:

1. Explicitly evaluate the ecological conditions required for acoustic interference, including assessing sustained spatiotemporal overlap, persistent masking, and the extent to which alternative niche axes reduce opportunities for acoustic interference.
2. Consider alternative mechanisms generating signal differentiation. Because signal divergence may emerge through multiple non-mutually exclusive mechanisms, including ecologically driven divergence and receiver-mediated processes, alone it provides limited support for acoustic interference-driven competition as the mechanism underlying acoustic space partitioning.
3. Integrate receiver perceptual systems into examinations of acoustic partitioning. Since acoustic interference emerges at the receiver, future studies should explicitly link signal differentiation to masking, detection, and discrimination processes.

By integrating ecological context, receiver perceptual systems, and evolutionary perspectives, future studies can better assess the contribution of competition for acoustic space to signal differentiation in mixed-species choruses.

