## Supplementary_materials for "A synthesis of the evidence for the acoustic space partitioning hypothesis"

**Electronic Supplementary Materials for the review: A synthesis of the evidence for the acoustic space partitioning hypothesis**

| Glossary | |
|---|---|
| Acoustic niche | Following Hutchinson's concept of the ecological niche [27], we define the acoustic niche as all conditions defining the signaling behavior of a species. This multidimensional space can be organized into three dimensions: a spatial dimension (where individuals signal), a temporal dimension (when individuals signal), and the acoustic space (the structure of the signal) (figure 1A). |
| Acoustic space | The acoustic space is an abstract multidimensional space defined by the acoustic features that characterize the signals of individuals within and/or among species in the chorus [28]. |
| Acoustic interference | The interaction of two or more sound waves producing a new wave, which is characterized by distinct acoustic characteristics, including frequency, amplitude, and phase. |
| Auditory masking | The sensory process by which the transduction of one sound alters the perception of another, resulting in reduced detection, localization, discrimination, and recognition of the signal. |
| Co-signalers | Two or more heterospecifics that signal at the same time and space and are part of the mixed choruses. |
| Mixed-species choruses | An aggregation of signalers of two or more species that share an acoustic signaling environment by overlapping in time (diel and/or seasonal) and space (habitat). |
| Focal mixed-species group | A set of species selected by researchers to address the questions assessed in their study. |

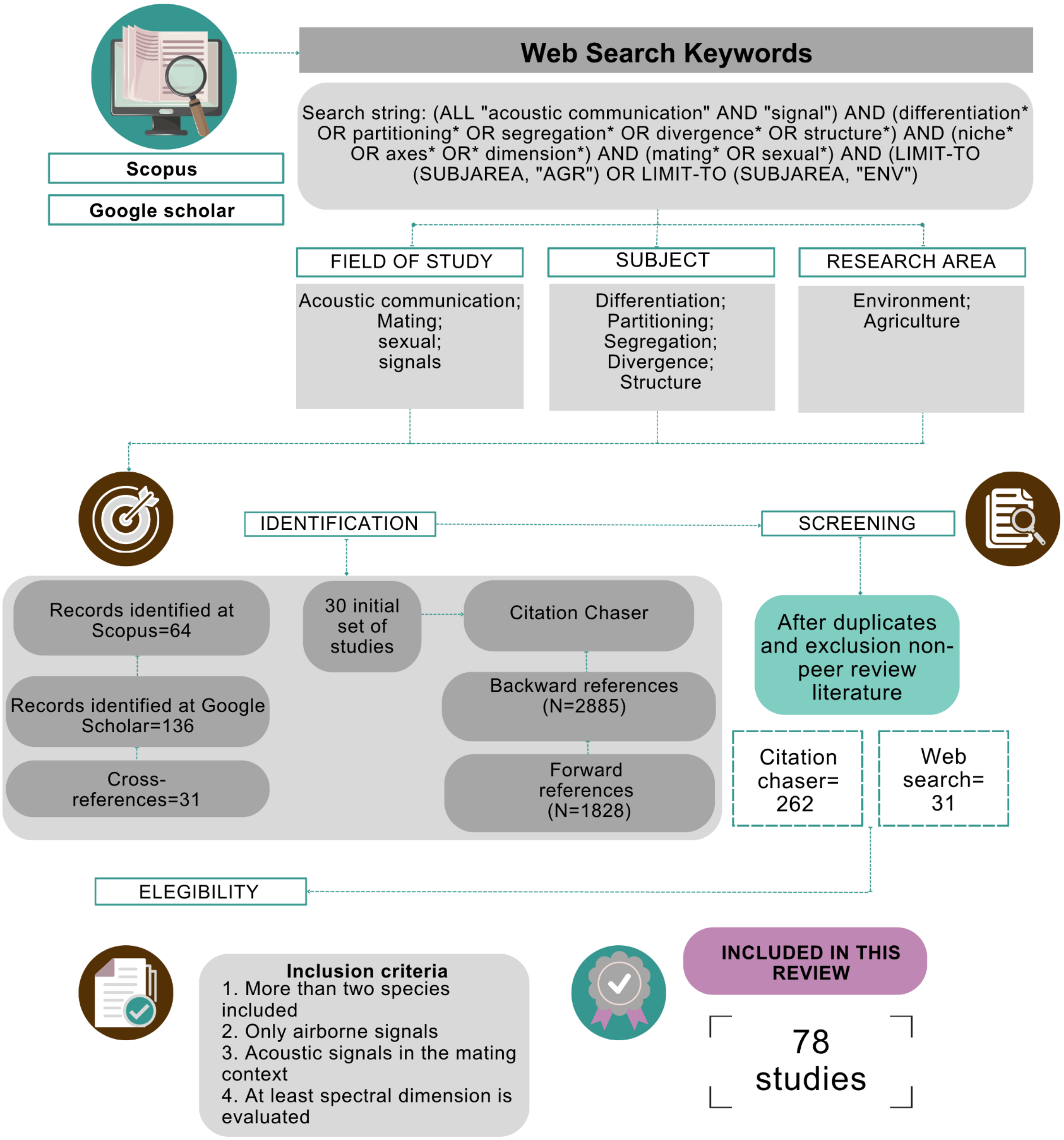


Fig. S1. PRISMA flowchart (Preferred Reporting Items for Systematic Reviews and Meta-Analyses) for the systematic review evaluating the acoustic space partitioning hypothesis in mixed-species groups. At the top, we indicate the two search engines used in our search, *Scopus* and *Google Scholar,* and list the words used per field of study, subject, and research area. We also describe the identification and eligibility criteria that led to the inclusion of the final 78 studies in our review.

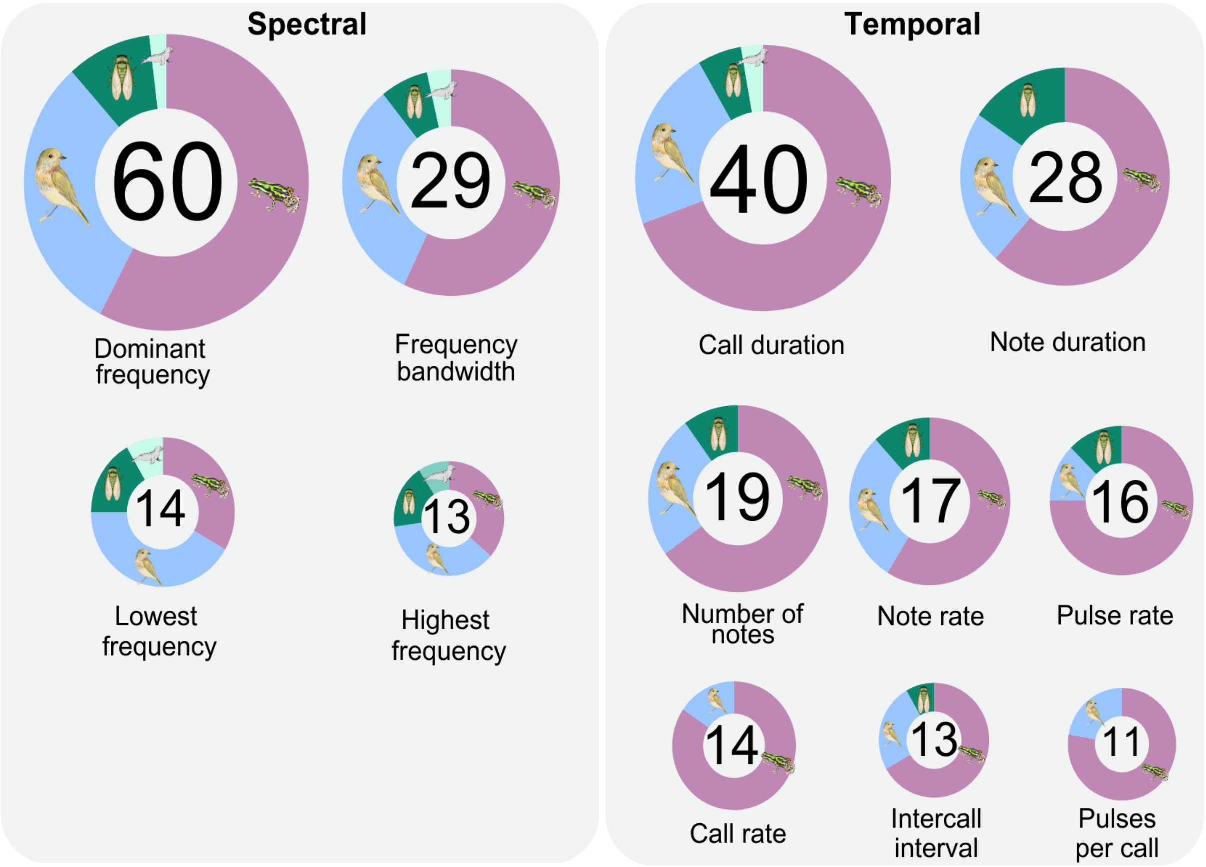


Figure S2. Most common acoustic features used to characterize signals in studies examining partitioning of the acoustic space across taxonomic groups. The numbers inside the ring represent the number of publications that evaluated each acoustic feature. Dominant frequency is the most commonly used feature, followed by call duration and frequency bandwidth. The colors represent the following taxa: purple = anurans, blue = birds, dark green = insects, and aqua = marine mammals.

| **Table S1.** Summary of acoustic space definitions extracted from 10 out of the 85 studies included in this review | | | |
|---|---|---|---|
| **Definition** | **Year** | **Taxa** | **Reference** |
| "each creature appears to have its own sonic niche (channel, or space) in the frequency spectrum and/or time slot occupied by no other at that particular moment" | 1987 | NA | Krause BL. 1987 The Niche Hypothesis: How animals taught us to dance and sing. *Whole Earth Review*, 6.<br><br>*not included in the systematic review |
| "acoustic space may be viewed as a limited resource that organisms compete for. Acoustic space is shared by all animals in a community but can only be partitioned in 2 primary dimensions (spectral and temporal)" | 2015 | Birds | Hart PJ, Hall R, Ray W, Beck A, Zook J. 2015 Cicadas impact bird communication in a noisy tropical rainforest. *Behavioral Ecology* **26**, 839–842. (doi:10.1093/beheco/arv018) |
| "The acoustic space can be represented as a resource that is shared by singing, calling, or stridulating animals" | 2016 | Insects | Roca IT, Proulx R. 2016 Acoustic assessment of species richness and assembly rules in ensiferan ommunities from temperate ecosystems. *Ecology* **97**, 116–123. (doi:10.1890/15-0290.1) |
| "The multidimensional channel through which species send and receive acoustic signals. Acoustic space is, therefore, a limited resource" | 2016 | Anura | Tennessen JB, Parks SE, Tennessen TP, Langkilde T. 2016 Raising a racket: invasive species compete acoustically with native treefrogs. *Animal Behaviour* **114**, 53–61. (doi:10.1016/j.anbehav.2016.01.021) |
| "…limited resource that is used by all vocal species in a community for species recognition, mate choice and resource defense, but the frequencies used are restricted by body size, phylogeny, habitat structure an biotic and abiotic sounds in the habitat" | 2017 | Anura, Birds, Insects | Aide TM, Hernández-Serna A, Campos-Cerqueira M, Acevedo-Charry O, Deichmann J. 2017 Species richness (of insects) drives the use of acoustic space in |

| | | | |
|---|---|---|---|
| | | | the tropics. *Remote Sensing* **9**, 1096. (doi:10.3390/rs9111096) |
| "Acoustic space represents an environmental resource that can be used by different species or lineages at the same time" | 2020 | Anura | de Lira AB, Alquezar RD, Giozza AP, dos Santos Campos ER, Cardoso AGT, Brandão RA. 2020 Acoustic space partition by four Hylidae in an open habitat in Central Brazil. *Heringeriana* **14**, 65–80. (doi:10.17648/heringeriana.v14i2.917916) |
| "ecological resource for which species compete and exclude one another" | 2021 | Anura | Allen-Ankins S, Schwarzkopf L. 2021 Spectral overlap and temporal avoidance in a tropical savannah frog community. *Animal Behaviour* **180**, 1–11. (doi:10.1016/j.anbehav.2021.07.024) |
| "…available resource used to recognize and discriminate between individuals of the same species or different species, leads to selective mating or non-selective between individuals from potentially divergent lineages"<br><br>"recurso disponible para reconocer y discriminar entre individuos de la misma u otras especies, que lleva al apareamiento selectivo o no-selectivo entre individuos de linajes evolutivos potencialmente divergentes" | 2021 | Anura | Ochoa-Ochoa LM, Ortiz-Ramirez MF, Figueroa-Huitrón R, Ríos-Muñóz CA. 2021 Ausencia de partición del nicho acústico en una comunidad de anuros en Chiapas, México. *ECOS* **30**, 1962. (doi:10.7818/ECOS.1962) |
| "…predictable distributions of acoustic traits in communities" | 2021 | Anura | Sugai LSM, Llusia D, Siqueira T, Silva TSF. 2021 Revisiting the drivers of acoustic similarities in tropical anuran assemblages. |

| | | | |
|---|---|---|---|
| | | | *Ecology* **102**, e03380. (doi:10.1002/ecy.3380) |
| "the acoustic space can become a limited resource since calling males need to deal with competitive pressure while receivers need to decipher messages in a noisy environment" | 2022 | Anura | Manzano MCR, Sawaya RJ. 2022 Environmental and phylogenetic aspects affect in different ways the acoustic niche of a frog community in southeastern Brazil. *An. Acad. Bras. Ciênc.* **94**, e20200705. (doi:10.1590/0001-3765202220200705) |
| "Acoustic space can be viewed as a niche axis that can be partitioned to avoid negative impacts of co-occurring signals" | 2022 | Anura | Allen-Ankins S, Schwarzkopf L. 2022 Using citizen science to test for acoustic niche partitioning in frogs. *Sci Rep* **12**, 2447. (doi:10.1038/s41598-022-06396-0) |

*Not included in the systematic review.

| **Table S2.** Number of mixed-species aggregations reported in studies of partitioning the acoustic space | |
|---|---|
| **Criteria Reported** | **Studies** |
| Number of species for one aggregation. | Azar & Bell (2016)<br>Balakrishnan et al. (2014)<br>Bignotte-Giró et al. (2019)<br>Blair (1958)<br>Bolaños-Sittler et al. (2021)<br>Bourne & York (2001)<br>Braga et al. (2023)<br>Chek et al. (2003)*<br>Chen et al. (2020)<br>Chitnis et al. (2020)<br>Chronister et al. (2023)<br>de Lira et al. (2020)<br>Diwakar & Balakrishnan (2007)<br>dos Santos et al. (2007)<br>dos Santos Protázio et al. (2015)<br>Duellman (1967)<br>Fouquette (1960)<br>Garcia-Rutledge & Narins (2001)<br>Gerhardt (1974)<br>Hart et al. (2015)<br>Heyer et al. (1990)<br>Hödl (1977)<br>Jain et al. (2014)<br>Knapton (1987)<br>Krishnan (2019)<br>Krishnan & Tamma (2016)<br>Latimer & Broughton (1984)<br>Lima et al. (2019)<br>Luddecke et al. (2000)<br>Luther (2009)<br>Malavasi & Farina (2013)<br>Manzano & Sawaya (2022)<br>Marquez et al. (1993)<br>Medeiros et al. (2017)<br>Mei et al. (2023)<br>Mossbridge & Thomas (1999)<br>Ochoa-Ochoa et al. (2021)<br>Planqué & Slabbekoorn (2008)<br>Pombal (2010)<br>Prasad et al. (2022);<br>Ríos-Lopez & Villanueva-Rivera (2013) |

| | |
|---|---|
| | Rossa-Feres & Jim (2001)<br>Schmidt et al. (2015)<br>Silva et al. (2008)<br>Sinsch et al. (2012)<br>Stanley et al. (2016)<br>Sueur (2002)<br>Sueur et al. 2010<br>Sugai et al. (2021)<br>Suzuki et al. (2012)<br>Tárano (2010)<br>Tennessen et al. (2016)<br>Tiwari & Diwakar (2019)<br>Tobias et al. (2014)<br>Torres et al. (2020)<br>van der Mescht et al. (2022)<br>Van Opzeeland et al. (2010)<br>Vasconcelos & Rossa-Feres (2008) |
| Number of species for more than one aggregation that are in the same area | Allen-Ankins & Schwarzkopf (2021)<br>Arriaga-Jaramillo et al. (2021)<br>Drewry GE & Rand AS. (1983)<br>Duellman & Pyles (1983)<br>Martins et al. (2006)<br>Salas et al. (1998)<br>Zimmerman (1983) |
| Number of species for more than one aggregation in different areas. | Cardoso & Price (2010)<br>Hart et al. (2021)<br>Kleyn et al. (2021)<br>Marín-Gómez et al. (2020)<br>Roca & Proulx (2016)<br>Shieh et al. (2015) |
| Number of species among multiple aggregations at different geographic areas. | Allen-Ankins & Schwarzkopf (2022)<br>Freitas & Toledo (2021)<br>Guerra et al. (2011)<br>Robert et al. (2019)<br>Villanueva-Rivera (2014) |

*Chek et al. (2003) provided information on several studies included later in this review; however, they reported one aggregation from their data.

| Table S3. Categories used to group studies assessing signal partitioning in mixed-species aggregations based on the statistical approach implemented | | |
|---|---|---|
| **Statistical approach** | **Description** | **Reference** |
| Observational | Acoustic features of each species signals are described verbally. Frequently, acoustic features are visually extracted from spectrograms and later compared between species. | Bignotte-Giró et al. (2019)<br>Blair (1958)<br>Duellman (1967)<br>Fouquette (1960)<br>Heyer et al. (1990)<br>Knapton (1987)<br>Luddecke et al. (2000)<br>Pombal (2010)<br>Ríos-Lopez & Villanueva-Rivera (2013)<br>Silva et al. (2008)<br>Van Opzeeland et al. (2010) |
| Basic statistics | Summarize the statistics that quantify the acoustic features of signals from different species. This approach usually provides a measure of central tendency (e.g., mean or median) and variation (standard deviation or standard error) for the distribution of acoustic parameters across signals from different species. | Bolaños-Sittler et al. (2021)<br>Chen et al. (2020)***<br>Diwakar & Balakrishnan (2007)<br>Drewry GE & Rand (1983)<br>Garcia-Rutledge & Narins (2001)<br>Gerhardt (1974)<br>Guerra et al. (2011)**<br>Hödl (1977)<br>Latimer & Broughton (1984)<br>Salas et al. (1998)<br>Sueur. (2002)<br>van der Mescht et al. (2022)<br>Villanueva-Rivera (2014) |
| Exploratory | Acoustic features are reduced to fewer dimensions to describe patterns within the dataset of species acoustic features in a mixed chorus. Non-linear Multidimensional Scaling analysis (NMDS), Principal component analysis (PCA), and discriminant | Allen-Ankins & Schwarzkopf (2022)<br>Allen-Ankins & Schwarzkopf (2022)**<br>Arriaga-Jaramillo et al. (2021)<br>Azar & Bell (2016)***<br>Bourne & York (2001)**<br>Cardoso & Price (2010)<br>Chen et al. (2020)***<br>Chitnis et al. (2020); |

| | | |
|---|---|---|
| | function analysis (DFA) are often used. | Duellman & Pyles (1983)**<br>Guerra et al. (2011)**<br>Kleyn et al. (2021)**<br>Krishnan (2019)***<br>Luther (2009)**<br>Mei et al. (2023)**<br>Prasad et al. (2022)**<br>Shieh et al. (2015)<br>Sugai et al. (2021)<br>Tárano (2010)**<br>Tiwari & Diwakar (2019)**<br>Tobias et al. (2014)** |
| Overdispersion | Assess the space between two points in a multi-dimensional space. For instance, it calculates the proximity of signals between the centroids of signals of different species to assess similarity. Examples of this approach include Euclidean distance and related measures, as well as Nearest Neighbor Analysis (NNA). Acoustic distance matrices are also included. | Allen-Ankins & Schwarzkopf (2022)**<br>Azar & Bell (2016)***<br>Chek et al. (2003);<br>dos Santos, & Rossa-Feres (2007)<br>Freitas & Toledo (2021)<br>Kleyn et al. (2021)**<br>Krishnan (2019)***<br>Lima et al. (2019)<br>Luther (2009)**<br>Manzano & Sawaya (2022)<br>Márquez et al. (1993)<br>Ochoa-Ochoa et al. (2021)<br>Schmidt et al. (2015)<br>Tobias et al. (2014)**<br>Vasconcelos & Rossa-Feres (2008) |
| Comparison among groups | Compare two distributions and determine if they are derived from the same underlying distribution. Parametric or non-parametric tests might be used to examine statistical differences among acoustic features of species signals.<br>Statistical examinations using t-tests or ANOVAs are common approaches. Contingency tables and Chi-square are also included. | Azar & Bell (2016)***<br>Bourne & York (2001)**<br>Braga et al. (2023)**<br>Chen et al. (2020)***<br>Duellman & Pyles (1983)**<br>Hart et al. (2015)<br>Hart et al. (2021)<br>Jain et al. (2014)<br>Krishnan& Tamma (2016)<br>Krishnan (2019)***<br>Malavasi & Farina (2013)<br>Martins et al. (2006)<br>Mei et al. (2023)** |

| | | |
|---|---|---|
| | Common distributions include Kolmogorov-Smirnov and Chi-squared. | Mossbridge & Thomas (1999)<br>Planqué & Slabbekoorn (2008)<br>Prasad et al. (2022)**<br>Tárano (2010)**<br>Tiwari & Diwakar (2019)**<br>Torres et al. (2020)<br>Zimmerman (1983)<br>Aide et al. (2017)<br>Allen-Ankins & Schwarzkopf (2021)<br>Balakrishnan et al. (2014)<br>Chronister et al. (2023)<br>Medeiros et al. (2017)<br>Robert et al. (2019);<br>Roca & Proulx (2016)**<br>Stanley et al. (2016)<br>Tennessen et al. (2016) |
| Indexes | Evaluates the partitioning of the acoustic space by creating categories based on the distribution of acoustic traits in the mixed chorus. Provides a measure of niche breadth based on the degree of overlap of species. Often referred to as the inverse of Simpson's diversity index (Pianka 1986) and Czechanowski index. | Braga et al. (2023)<br>de Lira et al. (2020)<br>dos Santos Protázio et al. (2015)<br>Marín-Gómez et al. (2020)<br>Roca & Proulx (2016)**<br>Rossa-Feres. & Jim (2001)<br>Schmidt et al. (2013)<br>Sinsch et al. (2012)<br>Sueur et al. 2010 |
| Predictive | Examines the relationship between a response variable, such as acoustic features, and one or more explanatory variables. This approach also allows for the study of interaction effects between variables that account for correlated observations. Some of these statistical analyses also allow individual predictions. Examples include Pearson's correlation analysis and regression, GLM, GLMM, LMM, and two-way interaction analysis. | Aide et al. (2017)<br>Allen-Ankins & Schwarzkopf (2021)<br>Balakrishnan et al. (2014)<br>Chronister et al. (2023)<br>Medeiros et al. (2017)<br>Robert et al. (2019);<br>Roca & Proulx (2016)**<br>Stanley et al. (2016)<br>Tennessen et al. (2016) |

| | | |
|---|---|---|
| Algorithms | Developed simulations to predict calling behavior using machine learning algorithms. | Suzuki et al. (2012) |

*The number of stars represents the number of statistical approaches used in the study

Complete list of studies included in this review

25. GUERRA, C., BALDO, D., ROSSET, S., BORTEIRO, C. & KOLENC, F. (2011) Advertisement and release calls in Neotropical toads of the Rhinella granulosa group and evidence of

natural hybridization between R. bergi and R. major (Anura: Bufonidae). *Zootaxa* **3092**, 26–42.
